# Audio-visual interactions in egocentric distance perception: Ventriloquism effect and aftereffect

**DOI:** 10.1101/2020.08.22.262444

**Authors:** Ľuboš Hládek, Aaron R Seitz, Norbert Kopčo

## Abstract

The processes of audio-visual integration and of visually-guided re-calibration of auditory distance perception are not well understood. Here, the ventriloquism effect (VE) and aftereffect (VAE) were used to study these processes in a real reverberant environment. Auditory and audio-visual (AV) stimuli were presented, in interleaved trials, over a range of distances from 0.7 to 2.04 m in front of the listener, whose task was to judge the distance of auditory stimuli or of the auditory components of AV stimuli. The relative location of the visual and auditory components of AV stimuli was fixed within a session such that the visual component was presented from distance 30% closer (V-closer) than the auditory component, 30% farther (V-farther), or aligned (V-aligned). The study examined the strength of VE and VAE as a function of the reference distance and of the direction of the visual component displacement, and the temporal profile of the build-up/break-down of these effects. All observed effects were approximately independent of target distance when expressed in logarithmic units. The VE strength, measured in the AV trials, was roughly constant for both directions of visual-component displacement such that, on average, the responses shifted in the direction of the visual component by 72% of the audio-visual disparity. The VAE strength, measured on the interleaved auditory-only trials, was stronger in the V-farther than the V-closer condition (44% vs. 31% of the audio-visual disparity, respectively). The VAE persisted to post-adaptation auditory-only blocks of trials, however it was weaker and the V-farther/V-closer asymmetry was reduced. The rates of build-up/break-down of the VAE were also asymmetrical, with slower adaptation in the V-closer condition. These results suggest that, on a logarithmic scale, the AV distance integration is symmetrical, independent of the direction of induced shift, while the visually-induced auditory distance re-callibration is asymmetrical, stronger and faster when evoked by more distant visual stimuli.

## 1 Introduction

The human ability to localize sounds requires a lot of flexibility. For example, whenever we enter a new room, we need to adapt the mapping from the acoustic localization cues to the perceived locations, to reflect the reverberation-related changes in the acoustic cues (Shinn-Cunningham, Kopco, & Martin, 2005). Vision plays an important role in these calibration processes, causing both an immediate re-alignment of the auditory spatial percepts to match the corresponding visual signals, and a long-lasting re-callibration of the acoustic localization cue mapping based on the visual inputs. The immediate alignment of auditory percepts with visual signals is known as the visual capture or the ventriloquism effect (VE; Jack and Thurlow 1973; Bertelson et al. 2000; Alais and Burr 2004; Recanzone 2009). Persisting visually induced re-callibration has been referred to as the ventriloquism aftereffect (VAE; (Kopčo et al., 2009; Recanzone, 1998; Wozny & Shams, 2011). VE and VAE have been extensively studied for horizontal sound localization. However, fewer studies have researched how vision influences auditory distance perception (Anderson & Zahorik, 2014; Calcagno, Abregú, Eguía, & Vergara, 2012; Cubick, Santurette, Laugesen, & Dau, 2015; Gardner, 1968; Mendonça, Mandelli, & Pulkki, 2016; Mershon, Desaulniers, & Amerson, 1980; Mershon, Desaulniers, Kiefer, Amerson, & Mills, 1981; Rébillat, Boutillon, Corteel, & Katz, 2012; Voss, 2016; Zahorik, 2001). The main goal of the current study is to systematically examine the basic properties of the ventriloquism effect and aftereffect in distance.

One previously reported property of ventriloquism in distance is that the strength of the effect depends on whether it is induced by visual stimuli displaced closer than the auditory signals (a condition refered to here as V-closer) vs. visual stimuli displaced farther away (a condition called here V-farther). For example, in anechoic space VE creates a strong illusion of the sound coming from the nearest plausible visual target located closer than the sound source (‘proximity image effect’; Gardner 1968; Min and Mershon 2005). Later, Mershon et al. (1980) found an effect similar to proximity image effect also for visual targets located farther than the sound source. These effects are strong in anechoic rooms in part because very few intensity-independent distance cues are available for these sources (Brungart & Rabinowitz, 1999). In reverberant rooms, the visual dominance in distance dimension can be expected to be weakened, as intensity-independent distance cues become available (Kopčo & Shinn-Cunningham, 2011). However, even in reverberation, auditory distance judgements are more accurate if the speakers are visible or if there are visual references in the room (Calcagno et al., 2012; Zahorik, 2001). In the current study, we examined whether there is an asymmetry in the strength of VE and VAE for the V-closer vs. V-farther stimuli in a reverberant room.

Most previous ventriloquism studies used one or only a few fixed visual stimulus locations combined with several auditory locations to examine the spatial dependence of the distance VE and/or VAE Zahorik (2003, Gardner 1968, Mershon et al., 1980). An important question that can elucidate the adaptive nature of the neural structures underlying ventriloquism is whether the auditory distance representation adaptaed is linear or non-linear (Bedford, 1993; Shinn-Cunningham, Durlach, & Held, 1998b). Typically, auditory distance studies analyze data on a logarithmic scale, as variance in distance judgments is approximately constant on this scale (Kopčo et al., 2012)(Anderson & Zahorik, 2014). Here we used a range of audio-visual stimulus pairs with a fixed A/V distance ratio to test whether adaptation by a constant amount in logarithmic units over a range of distances will induce constant VE and VAE shifts in logarithmic units. Such a result, as observed in horizontal adaptation studies (Shinn-Cunningham et al., 1998b), would indicate that the visually calibrated auditory distance representation is logarithmic, as opposed to, e.g., linear. We also wanted to examine the relative strength of the induced VAE vs. VE in both V-closer and V-furhter conditions, to determine whether the size of the aftereffect correlates with the size of VE and whether the adaptation caused by ventriloquism is equally large for the two directions. Finally, we were also interested in the dynamics of the ventriloquism buildup and decay, as VE and VAE occur on different time scales and the rates of adaptation might also be different for the V-closer vs. V-farther conditions.

To address these questions, a behavioral experiment was performed that investigated VE and VAE at intermediate egocentric distances of 0.7 to 2.04 m. The experiment was conducted in a small semi-reverberant room in which the targets were located directly in front of the participant. To present auditory (A) and visual (V) stimuli, we used an array of 8 linearly spaced loudspeakers and an array of LED lights densely covering the distances of 0.45 to 2.68 m, respectively. The ventriloquism effect was induced by presenting AV stimuli in which the A component (one of the 8 speakers) was paired with a V component (LED) positioned either 30% closer (V-closer) or 30% farther (V-farther). The size of V-component shift was fixed throughout the study so that the effects of displacement direction (V-closer vs. V-farther) and stimulus distance could be examined. The ventriloquism aftereffect was examined in A-only trials interleaved with the AV trials, and by separate A-only runs performed before and after adaptation. The interleaved A-only trials were expected to show the immediate aftereffect caused by preceding AV stimulation on the time scale of seconds, while the shifts observed in the separate A-only runs were expected to reflect a more persistent ventriloquism adaptation on the time scale of minutes. Each session took approximately one hour and consisted of several experimental runs in which the direction of V-stimulus displacement was fixed, combined with several pre-adaption and post-adaption control runs. While VE was expected to occur immediately upon the presentation of AV stimuli, VAE was expected to grow gradually as more and more AV stimuli were presented, and to decay gradually when the AV stimulus presentation stopped. The final analysis examined the VAE dynamics of adaptation across runs within a session, focusing on whether the VAE strength differs for the V-closer and V-farther conditions, which would be another indication that the adaptation process underlying VAE is direction-dependent.

Many previous auditory distance and ventriloquism studies used virtual acoustics to present stimuli. Here, the main goal is to understand the relative effects of visual stimuli on the auditory percepts, for which it is critical that the visual and auditory location percepts are veridical. Therefore, the study was performed in a real environment using loudspeakers and LEDs, to ascertain that the physical locations of the stimuli are unambiguhous and the percepts are as close to natural as possible. While this approach means that there are no issues related to virtual environment fidelity (e.g., issues with externalization or cross-modal binding), it also poses some limitations. The main ones are that 1) the range of A and V stimulus locations was limited by the room size, and 2) the sound received from individual loudspeakers varied slightly as loudspeakers cast an acoustic shadow affecting the more distant speakers when placed behind each other. However, these limitations are expected to only have small impact on the results of the study.

## 2 Methods

### 2.1 Participants

A group of 183 young adults participated in this study. All participants had normal or corrected-to-normal vision and normal hearing (by self-report). The experimental protocol was approved by the University of California, Riverside Human Research Review Board. All participants provided written informed consent.

### 2.2 Setup, stimuli, and procedures

The experiment was conducted in a small semi-reverberant acoustically treated room (broadband T60 = 408 ms; background noise 35 dB SPL) with internal dimensions of 2.6 m x 3.33 m x HEIGHT m (h). Participants were seated on a barber chair with adjustable height and a headrest. The chair back was facing the middle of one of the shorter walls in the room. The seated participant faced an array of nine uniformly spaced loudspeakers arranged one behind another (Fig. 1A) at the height of the listener’s ears. An acoustically transparent fabric covered the loudspeaker array, so the participant was unaware of the number and exact locations of the loudspeakers. Sound stimli were delivered from 8 loudspeakers (Peerless 830984, Tymphany) mounted on stands. An additional loudspeaker placed closest to the listener was only used to provide acoustic shadow. Visual stimuli were delivered from an array of fourty-eight uniformly spaced LEDs (4.76 cm spacing) mounted on a wooden frame raised 7 cm (at the far end) to 10 cm (at the near end) above the loudspeakers. The array height slightly decreased with distance so that all LEDs were clearly visible to the participants. The same LED array was used to collect responses. The loudspeakers and the LED array were coupled with a multichannel digital signal prcessor controller (RX-8, Tucker Davis Technologies, Alachua, FL, USA) and an 8-channel power amplifier (CROWN, Elkhart, IN, USA). A trackpad was used to collect the listener’s responses. An experimental computer outside the experimental room controlled the experiment, stimulus presentation, and response collection. The experimental procedure, stimulus generation and data analysis were implemented in MATLAB (Mathworks, Natick, MA, USA).

**Fig. 1.**
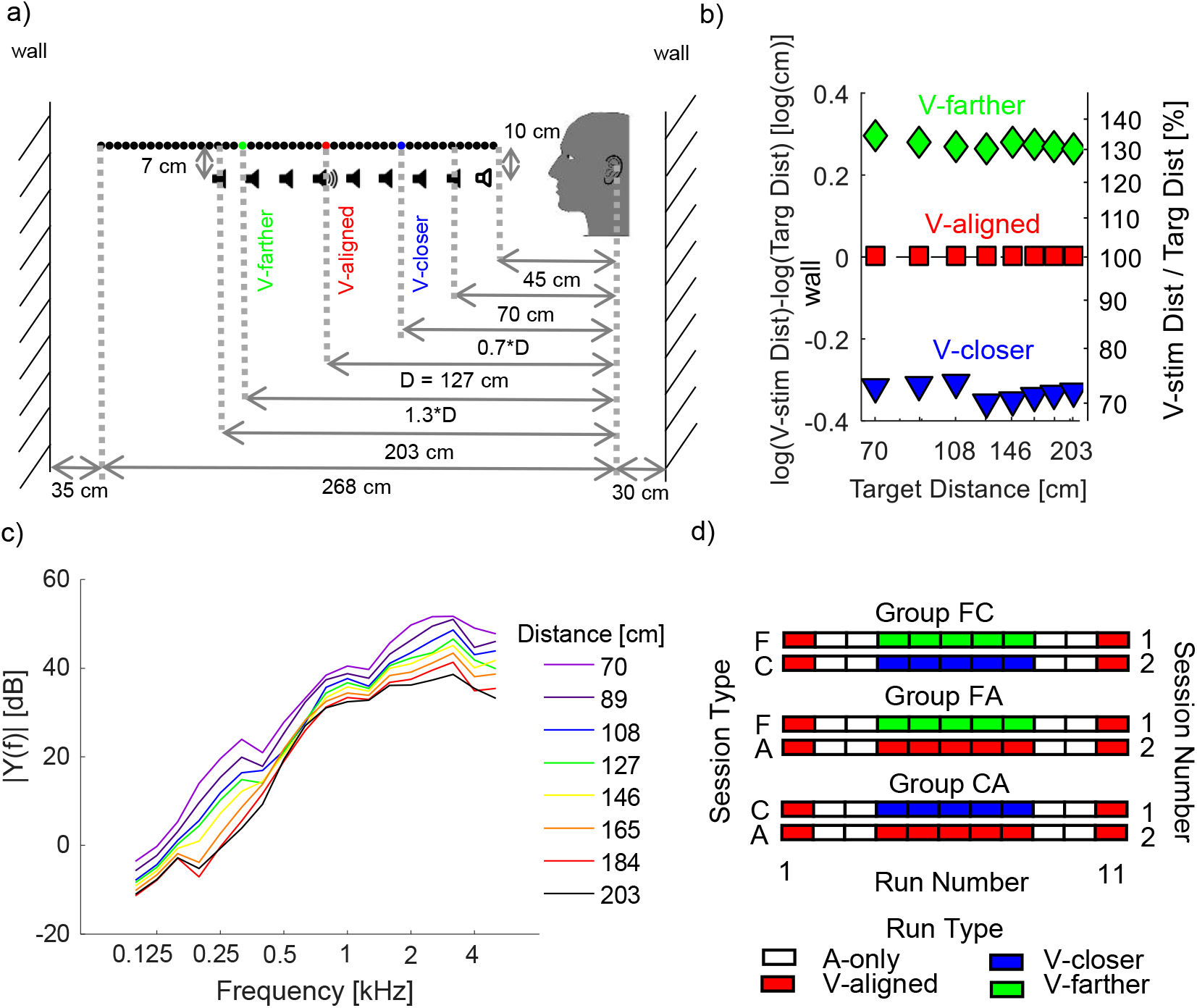
Setup, stimuli, and groups. (A) Setup and locations of the subject, loudspeakers, and LEDs in the room. The nearest loudspeaker (50 cm) was never used. An example AV stimulus pair is also shown, with the loudspeaker at D = 127 cm (shown as the active loudspeaker) and the corresponding V-farther LED (green), V-aligned LED (red), and and V-closer LED (blue). (B) Actual physical locations of the V-components of AV stimuli used to approximate the 30% AV separation in the V-farther and V-closer conditions. The V-component distance is shown re. the A-component (on a log scale) as a function of the target location. Right-hand side axis shows the ratio of distances in per cents. (C) Frequency response of the audio delivery system measured for each loudspeaker in 1/3-octave bands at the location of the listener head (without reverberation). (D) Schematic of the experimental sessions structure for different participant groups. Each participant was assigned into one of three groups and participanted in two sessions of eleven runs. Each row represents a series of runs performed by the participant from a given group from a given session. Sesssion types F, A, and C differed by the type of AV stimulus (V-farther, V-aligned, and V-closer) used in the adaptaion runs 4-8. Session ordering was counterbalanced across participants within each group.

The auditory stimuli were 300-ms broadband noise bursts. On each trial, one noise token was randomly selected from a set of 50 pre-generated tokens, and presented from one of 8 loudspeakers. The presentation sound level was held constant, which led to a natural decrease of sound level from 56 to 53 dB SPL with increasing target distance (measured, with reverberation, by a sound level meter Extech HD600, Flir Comercial System Inc., Elkhart, IN, USA). The frequency characteristics of the audio system and loudspeakers were estimated from impulse responses measured for each loudspeaker at the location of the listener’s head using the maximum length sequnce technique (Shinn-Cunningham et al., 2005). As shown in Fig. 1C, the responses are shifted systematically upwards as the distance of the speakers decreases. But, overall, the responses of the individual loudspeakers were aligned within ±3dB in 1/3-octave bands, when the distance-related gain change was accounted for. The visual stimuli consisted of 300 ms flashes of light from one of the LEDs. The audio-visual (AV) stimuli consisted of the auditory component (A-component, identical to the auditory stimulus) and the visual component (V-component, identical to visual stimulus) presented synchronously. The stimulus timing was controlled by a separate program run directly on the DSP. The V-component of the AV stimuli was either aligned with the A-component (V-aligned), placed approximately 30% closer than the auditory component (V-closer), or approximately 30% farther than the auditory component (V-farther). The exact relative placement of the V-componet vs. A-component deviated slightly from 30% because of the discrete positions of the LEDs (Fig. 1B). Also note that, while the 30% shift results in a constant shift as a function of target distance in log units when the V-closer and V-farther positions are considered separately, it corresponds to a larger absolute logarithmic value in the V-closer direction than in the V-farther direction. Specifically, 70% of target distance is −0.36 log(cm) while 130% distance is 0.26 (compare the left-hand vs. right-hand axis labels in Fig. 1B), which makes it more complicated to directly compare the effects of V-closer and V-farther adaptors in logarithmic units. We attempted to correct this disparity, as described in the results section.

Each participant participated in two experimental sessions, each consisting of 11 runs of 64 trials. Each trial started with the presentation of an A-only or AV-stimulus, followed by the participants’s response. The participant’s task was to indicate the perceived egocentric distance of the stimulus. When the AV stimulus was presented, participants were instructed to respond to the A component and ignore the V component. To collect the responses, a random LED was activated after the presentation of the stimulus. The participant then moved the active LED towards a perceived position using a trackball. Only one LED was active at any given time. The participant submitted their response with a button-click on the trackball (Wozny & Shams, 2011). No immediate feedback was provided, and after 500 ms inter-trial interval, the next trial was initiated. There were two types of runs. The A-only runs consisted of auditory-only trials. The AV runs consisted of A-only and AV trials pseudorandomly interleaved with the ratio of 3:1 (75% AV trials, 25% A trials). In each run, each of the eight target loudspeakers was randomly chosen to present stimulus on two auditory trials and six AV trilas. The relative displacement of the visual component was fixed within AV runs. So the AV runs were of 3 types, V-closer, V-farther or V-aligned. In each A-only run, each target loudspeaker was randomly chosen 8 times.

There were three types of sessions, F, C, and A, varying by what type of shift was being induced (V-farther, V-closer, or V-aligned, respectively; Fig. 1D). Each session, independent of its type, started with a pre-adaptation baseline period consisting of one V-aligned AV run followed by two A-only runs. The AV trials in run 1 were included both to minimize possible pre-adaptation biases that participants might have in their auditory-only distance responses and to establish a pre-adaptation reference. The adaptation period consisted of five AV runs (runs 4-8) with the AV disparity fixed within the session, which determined the type of session (C, F, or A). The post-adaptation period consisted of two A-only runs to assess the persistence of the induced recalibration and one V-aligned AV run to minimize any carry-over effects of adaptation on participant. The whole sessions took on average 60 mins to complete. There were 30-s breaks between the runs within a session. The participant received feedback about their progress during the resting breaks of the experimental sessions via voice commands played by one of the loudspeakers in the array.

Participants performed two sessions that were conducted on separate days. Each participant was randomly assigned to one of three experimental groups (Fig. 1D) differing by the conditions they performed in the sessions. Group CF (34 participants) performed the F and C sessions, Group FA (40 participants) the F and A sessions, and Group CA (40 participants) the C and A sessions. The order of sessions was counterbalanced across participants within a group.

A separate preliminary calibration measurement was performed to establish if any biases in the responses might be due to the experimental setup and, in particular, the response method based on a manually-controlled visual pointer and/or due to biases in visual distance perception. In the calibration measurement, a group of new 69 participants judged the egocentric distance of visual stimuli, using procedures identical to those described above, except for the following differences. The calibration session had only one run of 80 trials. On each trial a visual stimulus was presented from one randomly chosen LED in the array of 48 LEDs such that each LED was chosen at least once and not more than twice.

### 2.3 Data Analysis

The data from the sessions of the same type were pooled across the subject groups. E.g., the V-closer condition data were obtained from the C sessions of the participants from Group CF and Group CA. All responses were converted to the log scale (Anderson & Zahorik, 2014; Kopčo et al., 2012) before any manipulations. The graphs showing the results use the logarithmic scale, typically showing the difference between the logarithm of response distance and the logarithm of target distance on the primary y-axis and the ratio of the response distance and the target distance (in per cent) as the secondary y-axis (as already illustrated in Fig. 1B). Note, however, that the per cent scale cannot be used to describe the errorbars as the scale is non-linear. Unless specified otherwise, all results graphs show the across-subject mean and standard errors of the mean (SEM). Analysis of variance (ANOVA) with within-subject factor of distance (1-8) and between-subject factor of session type (F, C, A) assessed the statistical significance of the effects under the test. The statistical software CLEAVE (Herron, 2005) was used for the ANOVAs and the reported p values were corrected by using the Greenhouse-Geisser epsilon.

## 3 Results

The main goal of this study was to examine the effect of visual stimulation on auditory distance perception. To separate this effect from possible biases due to the experimental methods used here (like sensitivity to the exact placement of the speakers) and/or inherent biases in auditory distance perception, data from adaptation runs were analyzed relative to the pre-adaptation baseline runs 1-3. In this section, we first present the baseline data. Then, the effects of congruent and incongruent visual stimuli on the VE and VAE are presented as a function of target distance for the adaptation runs 4-8 and the post-adaptation runs 9-10. Finally, the dynamics of the VE and VAE build-up and breakdown are analyzed as a function individual runs within a session.

### 3.1 Baseline

Fig. 2 shows the baseline localization bias for the Visual-only stimuli (panel A), V-aligned AV stimuli (panel B), and Auditory-only stimuli (panel C). The visual baseline (Fig. 2A) was measured to validate the response method used here. The Visual-only responses were very accurate in the range in which the auditory targets were presented (70-203 cm), with only slight undershooting (by less than 5%) at the largest distances. However, the undershooting became more pronounced at larger distances, and reached 10% at the distance of 250 cm. It is not clear whether this bias is caused by actual biases of the visual distance perception or by some artifact of the the response method. However, it is likely that the response method influenced only the measurements for the most distant targets in the V-farther condition, for which the visual components were presented at distances larger than 2 m. To minimize the possible effect of this response method bias on the VA and VAE data, the data were corrected by adding the inverse of the response method bias to the baseline-corrected perceived location of auditory and AV targets (see below).

**Fig. 2.**
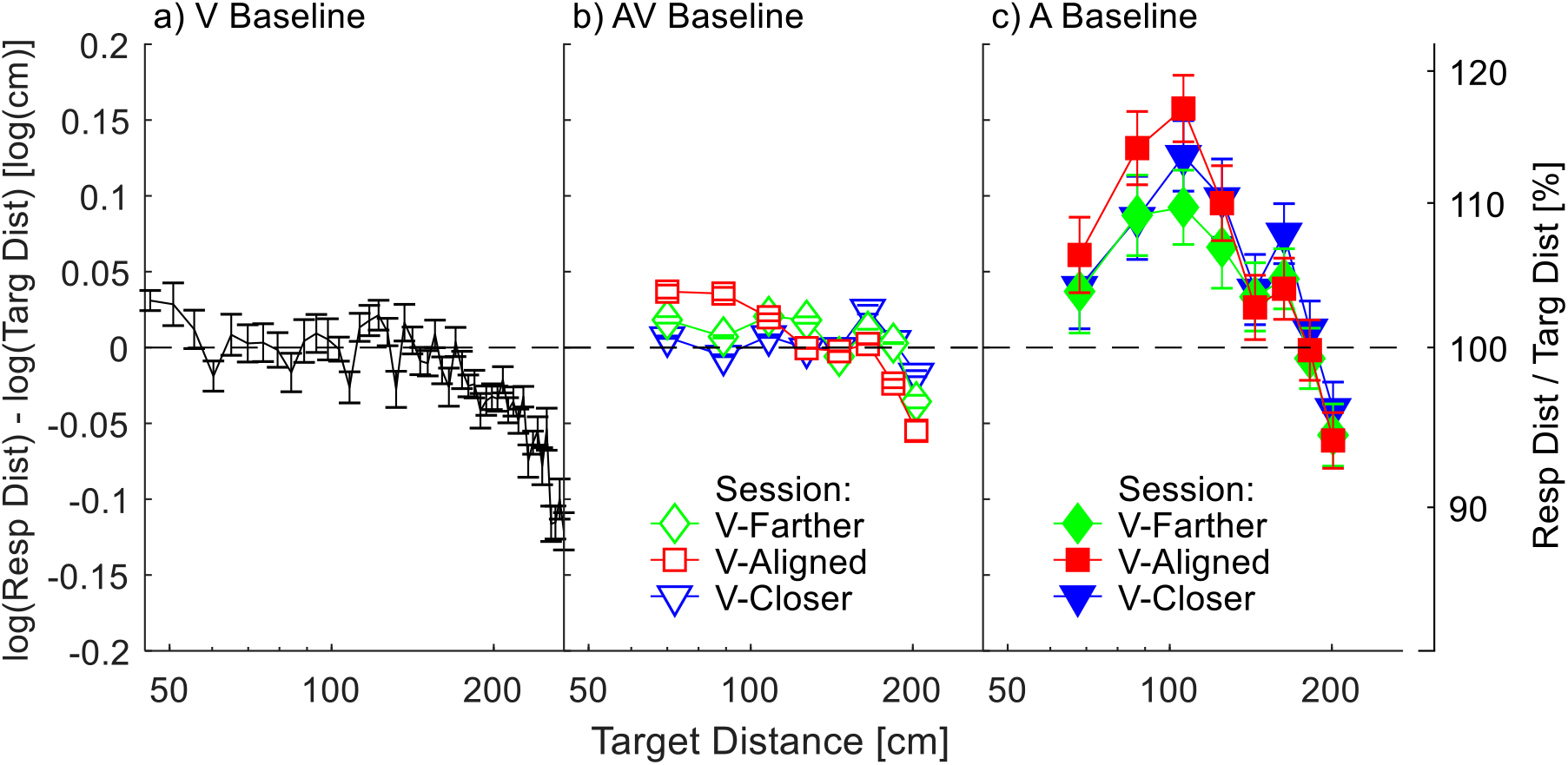
Baseline performance. Across-subject mean of the log of response distance *re*. the log of actual target distance as a function of the target distance. (A) Visual-only stimulus baseline measured in a calibration experiment. (B) Audio-visual baseline measured in run 1 of each session. (C) Auditory baseline from runs 1-3 in each session. Symbols in panels (B) and (C) represent data from three different sessions. The right-hand y-axis shows the ratio of the response and target distances in per cent.

The AV baseline (panel B) was measured in run 1 which used the V-aligned AV stimuli. Very little bias was observed (less than +/-3%), with an undershooting trend at the largest distances, consistent with the bias observed in the Visual-only calibration measurement (panel A). Importantly, there were no large differences between the three conditions (shown by different symbols), indicating that the subject groups were similar in their AV baseline performance.

Fig. 2C shows the auditory baseline obtained from auditory reponses in runs 1-3. Again, there were no systematic differences between subject groups (shown by different symbols). However, the participants tended to overestimate the auditory targets at most distances. The overestimation was largest (10-20%) at distances around 100 cm, and gradually decreased at larger distances, flipping to underestimation by approximately 5% at 200 cm (this underestimation, again, is likely caused by the measurement bias observed in Fig. 2A). This result is consistent with the auditory horizon observed in the previous studies which reported that distance estimates are overestimated for distances less than approximately 1-3 m and underestimated for distances larger than 1-3 m (Zahorik, 2001). In the current study, the location of the horizon appears to be at around 200 cm. Inconsistent with the auditory horizon explanation is the fact that the bias was reduced at the shortest distances. It is possible that this reduction is due to the presence of the interleaved AV stimuli in run 1, which provided subjects with information about the possible range of stimuli.

It may be somewhat surprising that such large biases in the A-only responses are observed, given that the A-only stimuli were interleaved with V-aligned AV stimuli for which no bias was observed (panel B). This result indicates that the V-component and A-component of the interleaved V-aligned AV stimulis might have been perceived at two separate locations. If that was the case, then it would be expected that ventriloquism aftereffect would gradually reduce the bias in the A-only responses during V-aligned adaptation due to multisensory enhancement (Burns et al, 2020). On the other hand, the bias might be caused mostly by the response method, e.g., due to the fact that responding was vision-based, and thus it might be easier to align a visual percept with a visually-based response than an auditory percept with a visually-based response. If that is the case, then no improvement due to VAE would be expected, as observed in the adaptation runs here (see section 3.2). Importantly, since this study focuses on relative effects of the V-farther and V-closer stimuli, it is assumed that these biases are not affecting the relative effects analyzed in the following sections.

### 3.2 Ventriloquism effect and aftereffect

The upper panels of Fig. 3 show the effect of visual stimulation on the perceived distance of AV (panel A) and A-only (panels B and C) stimuli with respect to the pre-adaptation baseline. The panels show the responses plotted in logarithmic units relative to the response distance baselines (from Fig. 2B-C). These data were corrected for the response method biases by adding the inverse of the response method bias (computed from data on Fig. 2A) to the baseline-corrected perceived location of auditory and AV targets. The data are plotted as a function of the actual target distance, separately for the three types of sessions (represented by different symbols and colors). Panel A shows the responses to AV stimuli during the adaptation runs 4-8, panel B shows the responses to the A-only stimuli interleaved with AV stimuli during the adaptation runs 4-8, and panel C shows the responses to the A-only stimuli in the A-only post-adaptation runs 9-10.

**Fig. 3.**
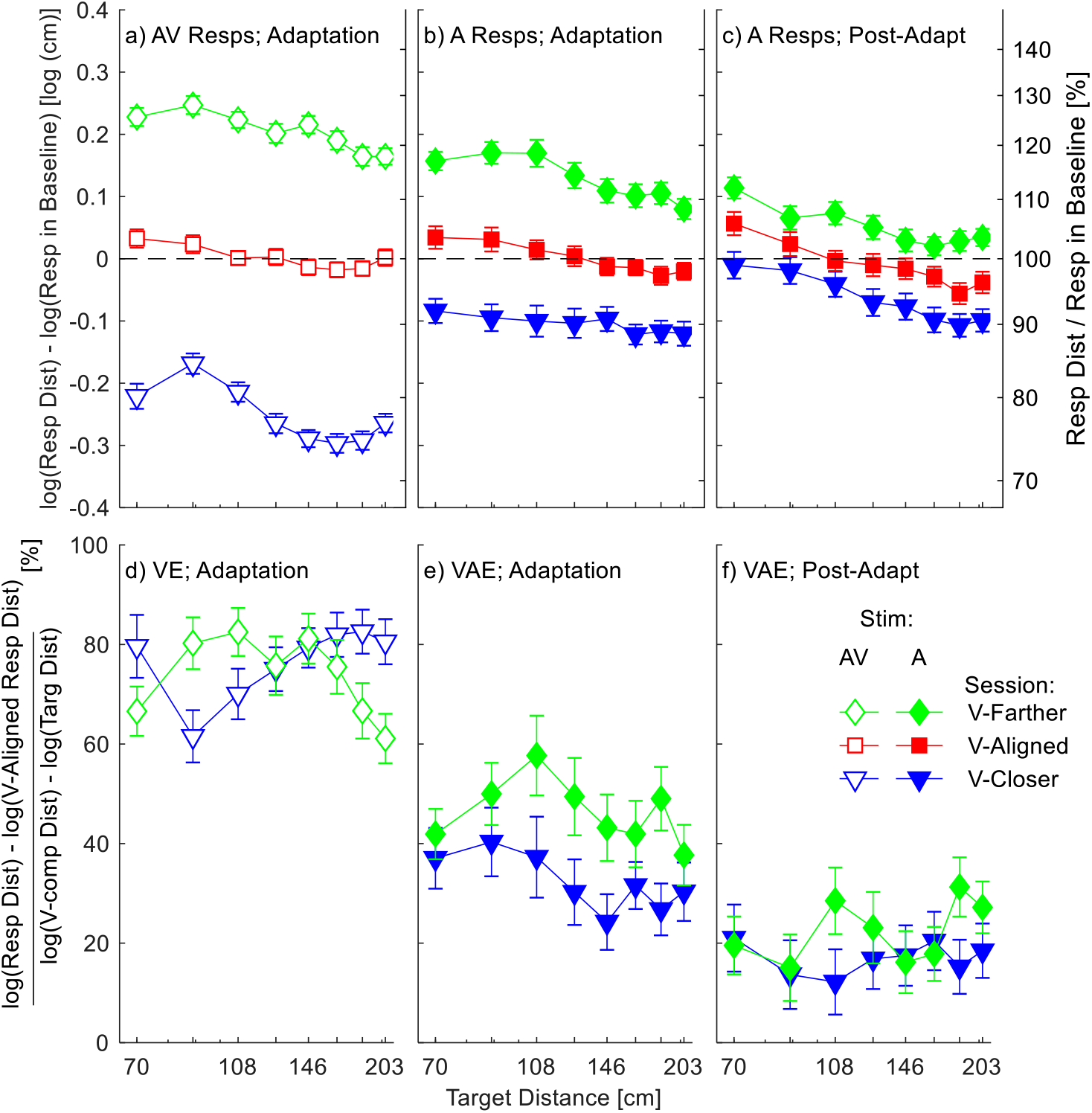
Performance in adaptation and post-adaptation runs. Panels A-C: Across-subject mean of the log of response distance *re*. the log of baseline-run response distance (from Fig. 2) plotted as a function of target distance. The three types of sessions are shown by different symbols. Panels A and B show the responses averaged across the adaptation runs 4-8 for AV targets (panel A) and A-only targets (panel B). Panel C shows the A-only responses averaged across the post-adaptation runs 9-10. The right-hand y-axis shows the ratio of the response distance and the baseline-run response distance (in per cent). Each of the lower panels (D-F) shows the V-closer and V-farther data from the corresponding upper panel, plotted relative to the corresponding V-aligned data (from the upper panel) and normalized by the physical disparity between the visual component and the target (from Fig. 1D; all in log units). All data were corrected for the response method bias shown in Fig. 2A.

The lower panels of Fig. 3 show the magnitudes of the ventriloquism effect (VE; panel D), immediate aftereffect (VAE; panel E), and persistent aftereffect (persistent VAE; panel F) derived from the data in the corresponding upper panels of the figure. Specifically, to compute the effect of a visual component placed closer (farther) than the auditory component in the lower panels, the differences between the corresponding V-closer (V-farther) and V-aligned data are taken from the upper panel and scaled by the physical disparities between the visual and auditory components (from Fig. 1D).

The AV responses in the V-aligned condition (red squres in Fig. 3A) are approximately at 0 log(cm), slightly overshooting for the near targets and undershooting for the distant targets. The V-farther (green diamonds) and V-closer (blue triangles) data are both shifted in the direction of the visual component, confirming the existence of the VE in the distance dimension. Also, their dependence on on the target distance is similar to that for the V-aligned data (slight downward slope). On average the V-farther responses are shifted up by 0.2 log(cm) and the V-closer responses are shifted down by 0.25 log(cm).

Similar to the AV-responses in panel A, the auditory-only responses during the adaptation runs (panel B) show a slight compression (downward slope) in all conditions. There is also a clear effect of the interleaved AV stimuli, such that both the V-closer (blue triangles) and the V-farther (green diamonds) stimuli shift the auditory-only responses in the direction of the visual component. The effect is approximately linear on the logarithmic scale (the blue and green lines are parallel with the red line), however it is weaker than in the AV responses (panel A), in particular for the V-closer data (compare the filled blue triangles of panel B to the open blue triangles of panel A).

In post-adaptation runs (Fig. 3C), the observed trends in the A responses are similar to those during adaptaion (panel B), but the magnitude of the effect of V-closer and V-farther adaptation is smaller than that in panel B, and approximately equal for all targets independent of their location or the direction of the visual component shift.

The downward sloping red graphs (as well as the other graphs) in Fig. 3A-C indicate that responses got more compressed during the adapation relative to the baseline. It is likely that this effect is a simple consequence of increased noise in subjects’ responses, either due to decreased attentiveness or increased tiredness during the experimental session. This bias can be mostly eliminated when comparing the effect of displaced V-stimuli (V-closer or V-farther) to the V-aligned reference. In summary, all of the results shown in Fig. 3A-C show a clear ventriloquism effect and aftereffect in distance perception. The effect appears approximately constant as a function of target distance and it depends in strength on the direction of induced shift (closer vs. farther). The following sections provide a detailed analysis of these results.

#### 3.2.1 Ventriloquism effect

Two trends in Fig. 3A suggest that additional corrections need to be applied before the effects of interest here can be evaluated. First, the AV response biases in Fig. 3A are overall larger for the V-closer stimuli than the V-farther stimuli (relative to the V-aligned stimuli; compare the blue triangles and green diamonds to the red squares). However, this asymmetry is likely caused by the fact that a 30% shift of the visual component corresponds to a larger disparity in logarithmic units for the V-closer than V-farther condition (compare the left-hand and right-hand y-axis in Fig. 3A-C). Second, while the V-farther data (green diamonds) are relatively smooth, the V-closer graph (blue triangles) has a peak at the 89-cm target and a trough at the 165-cm target. This non-unifromity might be due to the slight variability in the size of the physical disparity between the V and A components, which was not always exactly 30% (Fig. 1B). To account for these artifacts, as well as for the compressive bias observed even in the V-aligned condition, the V-closer and V-farther data from panel A were re-plotted in Fig. 3D relative to the across-subject average of the V-aligned data, and scaled by the size of the physical AV disparity. Thus to show the strength of the ventriloquism effect, Fig. 3D plots the size of the shift in the perceived location as a proportion of the shift in the physical displacement of the V-component which induced it.

VE data from Fig. 3D were subjected to a two-way mixed ANOVA with factors of target location and condition, which showed that the VE is approximately equally strong in the V-closer and V-farther conditions (main effect of condition: F(1,146)= 0.29, n.s.). While there was a considerable variation in the size of the effect as a function of target distance in both the V-closer and V-farther graphs (interaction of condition and target distance: F(7,1022)= 5.97; p<0.0001), there was no clear systematic trend of increasing or decreasing effect size with target distance. Moreover, this variation is relatively small compared to the overall VE and it is likely driven by noise, e.g., due to factors like the limited accuracy in the estimation of the V-aligned responses. Specifically, the observed pattern of maxima and minima in the V-closer data is exactly mirroring that in the V-farther data (i.e., the maxima in the green line are aligned with the minima in the blue line, and vice versa), which would be expected if the effect is mostly driven by noise in the V-aligned data estimation, as that estimate gets subtracted from the V-farther data and added to the V-closer data. Therefore, the dependence of the results on target distance was not further considered.

In summary, the effect of spatially displaced visual stimulus on the perceived distance of a simultaneously presented auditory stimulus is approximately constant on a logarithmic scale at 72% of the visual stimulus displacement when the visual component is shifted by a constant 30% *re*. auditory component over a range of distances of 0.7 to 2 m. This effect is independent of whether the visual component is located closer or farther than the auditory component (V-closer vs. V-farther) and approximately independent of the absolute distance of the auditory stimulus (ranging from 60 to 80%). Thus, there is no evidence that the effect measured in a room would be stronger for the V-closer direction, as suggested for anechoic space by Gardner (1968). And, the data provide further support for the suggestion that auditory distance is represented in logarithmic units, as the observed VE was approximately constant at 72% of the imposed 30% AV disparity in all the VE measurements when expressed in logarithmic units. Note, also, that this 72% value is considerably smaller than that observed for VE in the horizontal direction, in which the effect can reach more than 95% of the imposed disparity (Bruns, Liebnau, & Röder, 2011; Frissen, Vroomen, & de Gelder, 2012; Kopčo et al., 2009; Lewald, 2002; Recanzone, 1998, 2009; Stekelenburg, Vroomen, & de Gelder, 2004; Vroomen, Bertelson, & de Gelder, 2001).

#### 3.2.2 Ventriloquism aftereffect

The ventriloquism aftereffect (VAE) was examined on two time scales. First, the immediate VAE was assessed for the A-only trials randomly interleaved with the AV trials during the adaptation runs. This effect reflects adaptation on the time scale of seconds and tens of seconds. Second, persistent VAE was assessed by two post-adaptation A-only trial runs, reflecting adaptation on the time scale of minutes.

To evaluate the immediate VAE, data from panel B in Fig. 3 were replotted in panel E relative to the V-aligned data and scaled by the size of the physical AV disparity, using the same procedure and rationale as discussed above for the VE. These data were subjected to a mixed two-way ANOVA with the factors of target location and condition which showed that VAE was independent of the target location (F(7,1022)= 1.81; p=0.08), but that it was clearly stronger for the V-farther condition (approximately 44% of the AV offset) than for the V-closer condition (approximately 31% of the AV offset; F(1,146)= 5.48; p=0.02).

To evaluate the persistent VAE, the data from panel C were again rescalled and replotted in panel F. The observed patterns are also similar to the immediate VAE, except that the effect is smaller (on average, 19% of the physical AV displacement). ANOVA performed on these data did not show any significant effect or interaction (p > 0.1). However, the green diamond data tend to fall above the blue triangle data in panel F, showing that there still is a trend for the aftereffect to be stronger in the V-farther condition.

The ventriloquism aftereffect induced by the AV disparity is similar to the ventriloquism effect in that it is constant as a function of distance. However, contrary to the VE, the VAE is asymmetrical, considerably stronger in the V-farther direction. Speciffically, when the immediate VAE is expressed as a proportion of VE, it is on average 63% in the V-farther direction and only 43% in the V-closer direction. When the persistent VAE is expressed as a portion of VE, it is no average 29% and 22% in the V-farther and V-closer conditions, respectively. There are at least two possible explanations of this farther-closer asymmetry. Either the neural distance representation adapted in the ventriloquism aftereffect is more flexible in one direction than the other, or the visual signals that cause the adaptation are stronger or more salient in the V-farther direction. Importantly, the fact that VAE observed here is constant as a function of target distance for both directions suggests that the neural representation of auditory distance is adaptable in logarithmic units, at least over the range of distances and for the AV disparity examined in the current study.

Finally, no adaptation was observed in the V-aligned data, even though the V-aligned A-only baseline was shifted considerably re. the AV baseline in Fig. 2C vs. Fig. 2B. It would be expected that, if the baseline misalignment was perceptual, the shift would get corrected by the V-aligned signals during the VAE adaptation due to multisensory enhancement (Bruns, Dinse, & Röder, 2020). Since no correction occurred, the disparity in baselines is more likely caused by differences in response strategies than in perception. E.g., since the same LED array was used to present the V-stimuli and collect the responses, the responses might be more accurate for stimuli with V-components than for auditory-only stimuli, as the subjects might be able to directly compare the perceived location of the visual-stimulus and the response LED.

### 3.3 Build-up and break-down of the ventriloquism effect and aftereffect

No systematic effects of target distance were observed in the analysis in the preceding section. Therefore, the dynamics of the build-up and break-down of the visually-induced adaptation were analyzed after collapsing the data across target location (while keeping the data from each experimental run separate). It was expected that the VE build-up and break-down will be immediate while the VAE build-up and break-down would have slower dynamics. Also, it was expected that, given the difference in the strength of the VAE for the V-farther vs. the V-closer condition, there might be a difference in the rate at which the VAE builds up for the two directions.

Fig. 4A shows the localization biases in the AV (empty symbols) and A-only (full symbols) trials separately for each run as a function of the run number, referenced to the pre-adaptation runs 1-3. Fig. 4B shows the magnitudes of the VE and VAE computed by referencing the data from panel A to the V-aligned condition and scaling them by the physical disparity of the visual and auditory components (as in the previous section). Fig. 4C aims at estimating the average dynamics of the VAE in the V-closer vs. V-farther condition by showing the average of the VAE build-up (runs 4-5 referenced to the average of preadaptation runs 1-3 from panel B) and the inverse of the VAE break-down (runs 9-10 referenced to run 8 from panel B).

**Fig. 4.**
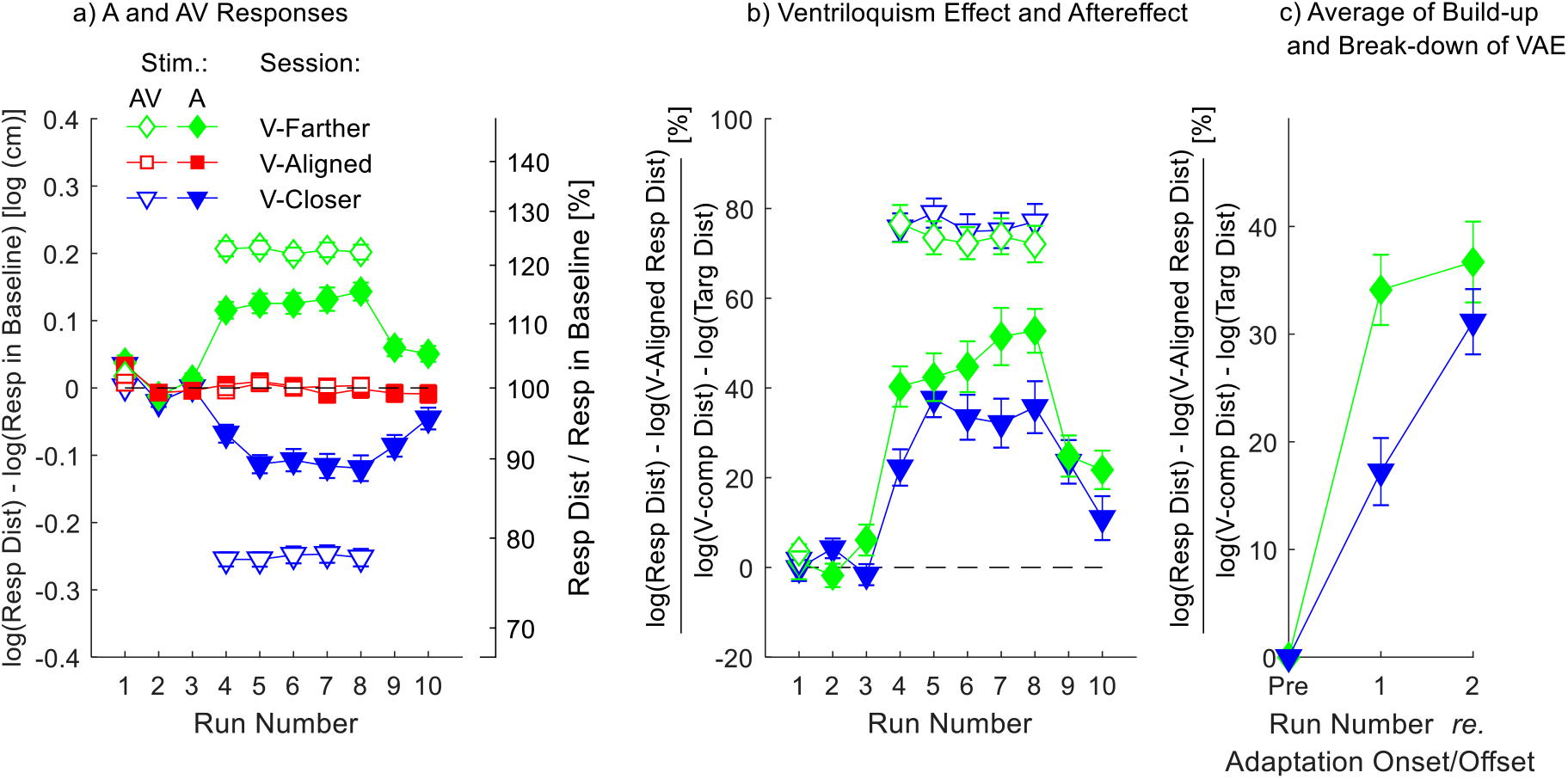
Temporal profile of ventriloquism adaptation. (A) Response biases as a function of the run number within a session for the three different conditions, averaged across target distance and referenced to the pre-adaptation baselines (runs 1-3). (B) VE (open symbols) and VAE (filled symbols) as a function run number, computed from data in panel A by referencing the V-closer and V-farther data to the V-aligned data and scaling them by the physical disparity between the stimuli. (C) Dynamics of post-onset/offset VAE adaptation shown as an average of the build-up (runs 4-5) and inverse of break-down (runs 9-10) of VAE from panel B, referenced to the final runs prior to the onset (runs 1-3) and offset (run 8) of the AV stimuli.

We can see in Fig. 4A that, averaged across target locations, the V-aligned data were stable near 0 log(cm) throughout the sessions (red open and filled diamonds are always near 0). The VE was fast and stable (open green diamonds and blue triangles show constant values throughout the adaptation runs 4-8 in Fig. 4A), while the VAE had a clear build-up and decay after the onset and offset of adaptation (full green diamonds and blue triangles grow gradually in runs 4-8 and decayg radualy in runs 9-10). When expressed as a proportion for physical disparity (Fig. 4B), the VE magnitudes were approximately equal for the V-closer and V-farther conditions, at 70-80% of the the V-component offset (open symbols). The VAE expressed as a proportion of physical disparity was slower than VE and had more complex dynamics. Confirming the results described in the previous section, VAE was stronger for the V-farther than V-closer conditions during both the adaptation and the post-adaptation runs (green filled diamonds are mostly above blue triangles for runs 4 – 10 in Fig. 4B). Importantly, the results also showed a difference in the rate of adaptation. The V-farther data (filled green diamonds in Fig. 4B) were approximately 40% of AV disparity by the first adaptation run (run 4), while the V-closer data (filled blue triangles in Fig. 4B) were only 22% of the AV disparity within the first adaptation run. Also, the V-farther data continued to grow, reaching more than 50% by run 8, while the V-closer data were approximately constant at around 35% in runs 5-10. A similar pattern was observed after the offset of the adaptation. V-farther data dropped much faster than the V-closer data (compare runs 9 vs. 8 for the filled green diamonds and blue triangles). To better visualize this difference in adaptation rate, Fig. 4C plots the average of the build-up and the inverse of the break-down data from the two runs post-onset and post-offset of adaptation (relative to the preceding runs). The panel clearly shows a much faster growth in the V-farther adaptation in the first run after its on/offset, while the V-closer adaptation grows much more gradually.

Statistical analysis confirmed these results. First, ANOVA performed on the VE data from Fig. 4B with the factors of run (4-8) and condition (V-closer vs. V-farther) found no main effect or interaction of the condition and run factors. A similar ANOVA performed on the VAE data from Fig. 4B only found a significant main effect of the condition (F(4,584)=4; p<0.01). Finally, ANOVA performed on data from Fig. 4C found a significant main effect of run (F(1,73)= 16.94; p<0.01), a significant main effect of condition (F(1,73)= 11.51;p<0.01), and a significant interaction of the two factors (F(1,73)= 10.22; p<0.01).

Overall, the difference in the dynamics of the VE vs. VAE supports the idea that these two effects are caused by two different mechanisms. The VE is fast because this effect is caused by immediate cross-modal integration of the distance information from the auditory and visual modalities. And, the current data show that the weight used for each modality in the integration is approximately constant even across multiple experimental runs. On the other hand, the VAE is slow because it entails neural adaptation of the perceptual map of auditory space guided by the visual signals. Moreover, the current results show that the neural adaptation is faster when the visual signals guide it to perceive the auditory targets as farther away than when they guide it to shift the percepts to closer.

## 4 Discussion

The current study examined ventriloquism effect (VA) and aftereffect (VAE) in the distance dimension for auditory stimuli presented over a range of distances of 0.7 – 2.03 m in real reverberant environment in front of the linstener. It found that, for a fixed 30% relative shift of the visual component of the AV stimuli, the induced ventriloquism effect is constant at approximately 72% of the V-displacement, roughly independent of the stimulus distance or the direction of the induced shift on the logarithmic scale. Also, this VE strength was constant over time. On the other hand, the ventriloquism aftereffect induced by these stimuli, while still independent of target distance, was stimulus-direction dependent. The VAE was stronger when the V-component of AV stimuli was placed farther than the auditory component (reaching 44% of the V-displacment) than when it was placed closer (31%). Also, the VAE in the V-farther condition was faster than in the V-closer condition.

The observed differences between VE and VAE confirm that these effect are caused by two different processes, comparable with the VE and VAE in the horizontal dimension (Bosen, Fleming, Allen, O‘Neill, & Paige, 2017; Bruns et al., 2011; Kopčo et al., 2009). Specifically, VE is likely a consequence of an immediate cross-modal integration, while VAE is a result of recalibration of some auditory spatial representation by the visual signals. For VE, the observed independence of the effect of the direction of shift is not consistent with the “proximity image effect” (Gardner, 1968) which showed that, in anechoic room, the ventriloquism effect is stronger in the V-closer direction. A possible explanation of this difference is that the current experiment was performed in a real reverberant room and the subjects were provided with a level-independent auditory distance cue, the direct-to-reverberant energy ratio, whereas in the Gardner study no such cue was available and the subjects likely relied on guessing. The observed constant 72% weight given to the V-component in the current study is much smaller than the weight typically observed in horizontal studies, in which this weight is commonly around 90% (e.g., Kopčo et al. 2019). Assuming that auditory and visual components were combined optimally (Alais & Burr, 2004) in the current study, this result suggests that the auditory distance acuity is much more comparable to the visual distance acuity than in the horizontal dimension. However, more studies need to be performed to determine whether the observed 72% V-component weight generalizes to other AV disparities, other reference distances, and other types of stimuli (e.g., familiar 3-dimensional objects might provide more accurate visual distance information than the single LEDs used here, while familiar sounds like speech stimuli might provide more auditory distance information).

Even though ventriloquism effect observed in this study was equally strong in the V-farther and V-closer conditions, the ventriloqusim aftereffect evoked by it was stronger in the V-farther condition than in the V-closer condition. This asymmetry in the effect was not expected and it is not immediately clear why it has occurred. It might be related to the “adjacency effect” reported in auditory distance perception by Min & Mershon (2005), which observed that manipulating the perceived distance of a nearby sound by a simultaneously presented distal light (i.e., ventriloquism effect) caused an adjacent reference sound (i.e., ventriloquism aftereffect) to be perceived as farther away, whereas the reverse setup did not cause an adjacent reference sound to be perceived as as closer. The current results suggest that this asymmetry is a general observation in the recalibration of auditory distance percepts by vision, and that it also applies to the speed with which the aftereffect builds up. Alternatively, this effect might relate to the fact that the A baseline responses were overestimated even in the preadaptaion in the current study, which means that the perceived A-only locations were much closer to the V-farther visual adaptors than V-closer visual adaptors, even when they were interleaved with V-aligned AV stimuli. Finally, the constant 30% disparity used in this study corresponds to a larger disparity in logarithmic units for the V-closer condition (−0.36) than for the V-farther condition (0.26) (Fig. 1B). In the analysis performed here we scaled all the observed effects by the actual applied separation (in log units), thus correcting for this difference. However, it is possible that the strength of AV binding was stronger in the V-farther condition, as the separation in log units was smaller, which might have resulted in the observed stronger adaptation. Future studies will need to independently control the physical and perceptual disparities in the auditory and visual components of the audiovisual and auditory stimuli in order to disentangle these alternatives.

The VAE data of the current study show that, in both V-closer and V-farther conditions, the participants adapt to a constant-ratio shift in visual-component distance over a range of distances by a shift in response that is, again, constant over the examined distance range. Previous studies in the horizontal plane suggest that adaptation in response to a non-linear transformation of space often leads to a linear response pattern that approximates the non-linear transformation (Shinn-Cunningham, Durlach, & Held, 1998a). Based on these results, one can assume that the neural structure undergoing adaptation in distance ventriloquism is also adapting linearly in the units in which distance is represented. I.e., the current results show that the ratio scale (or, equivalently, the logarithmic scale) is the scale used by the structure adapted by ventriloquism in distance. This result is also consistent with the variance in responses to auditory stimuli which is approximately constant across distances (Anderson & Zahorik, 2014; Kopčo et al., 2012). However, given that the ratio scale was the only scale examined here (i.e., we did not check whether the adaptation was equally good for a linearly constant shift in the V-componenet), it is also possible that the representation is flexible enough to be capable of adapting to several various transformations evoked by visual stimuli (in the horizontal domain, ventriloquism adaptation can be obtained both by shifting the V-component by constant amount or by varying the gain of the V-componenet displacement; Zwiers et al. (2003)), or that it is even robust to adaptation size that is varying from trial to trial (as was recently shown for horizontal ventriloquism; (Bruns et al., 2020)).

In terms of the temporal profile, the large immediate VAE and smaller persistent VAE observed here are generally consistent with the observations of multiple distinct cross-modal recalibration mechanisms reported in the horizontal VAE studies (Bosen, Fleming, Allen, O’Neill, & Paige, 2018; Watson, Akeroyd, Roach, & Webb, 2019). However, a signle adaptation mechanism cannot be ruled out for the current results given that the current study was not specifically designed to examine this question. On the other hand, the asymmetry in the adaptation rates between V-closer and V-further conditions observed here differs from horizontal ventriloquism in which no equivalent left-right asymmetries are observed. Future studies will need to examine whether the observed asymmetry in the adaptation rates is directly related to the observed difference in the strength of the induced VAE, or whether these two phenomena are separable.

Finally, an unexpected result of the current study is that A-only responses were overestimated in the preadaptation baseline measurements and in the V-aligned condition even though multisensory enhancement would be expected to occur, reducing the biases in the A-only stimuli (Anderson & Zahorik, 2014; Bruns et al., 2020). It is difficult to identify the cause of this disparity, as no A-only initial runs were performed to serve as a basline. Thus, it is for example possible that the enhancement has actually occurred and that the biases would have been much larger in the initial run if it was performed without any interleaved V-aligned AV stimuli. However, even if that was the case, it still is not clear why the bias would not continue to reduce to 0 even during the 5 adaptation runs in the V-aligned condition. Future studies are needed to explore this effect.

## 5 Conclusions

The current results show that ventriloquism effect (VE) and aftereffect (VAE) in the distance dimension can be robustly induced in real reverberant environment, and that they are approximately independent of target distance on a logarithmic scale. They also show that while VE is immediate and independent of the direction of induced shift, VAE is more complex, showing stronger and faster effect in the V-farther direction. Future studies need to explore the possible cause of this asymmetry and its implications for our understanding of the mechanisms of audio-visual integration in the human brain.

## Acknowledgements

This work was supported by EU H2020-MSCA-RISE-2015 Grant No. 691229, by UPJŠ VVGS-2020-1514, and by BCS-1057625 to AS.

